# Genome-wide CRISPR/Cas9 Screens Reveal Shared and Bespoke Mechanisms of Resistance to SHP2 inhibition

**DOI:** 10.1101/2022.08.26.505487

**Authors:** Wei Wei, Mitchell J. Geer, Xinyi Guo, Igor Dolgalev, Neville E. Sanjana, Benjamin G. Neel

**Affiliations:** Laura and Isaac Perlmutter Cancer Center, NYU Grossman School of Medicine, NYU Langone Health, New York, NY; Department of Biology, New York University, New York, NY; New York Genome Center, New York, NY

## Abstract

SHP2 (*PTPN11*) acts upstream of SOS1/2 to enable RAS activation. Allosteric inhibitors (SHP2is) stabilize SHP2 auto-inhibition, preventing activation by upstream stimuli. SHP2is block proliferation of RTK- or cycling RAS mutant-driven cancers and overcome adaptive resistance to other RAS-ERK pathway drugs. Several SHP2is are in clinical trials. To identify potential SHP2i resistance mechanisms, we performed genome-wide CRISPR/Cas9 knockout screens on two SHP2i-sensitive AML cell lines and recovered genes expected to cause resistance, including tumor suppressor (*NF1*, *PTEN*, *CDKN1B*) and “RASopathy” (*LZTR1*, *RASA2*) genes, and several novel targets (*INPPL1*, *MAP4K5,* epigenetic modifiers). We then screened 14 cancer lines with a focused CRISPR library targeting common “hits” from the genome-wide screens. *LZTR1* deletion conferred resistance in 12/14 lines, followed by *MAP4K5* (8/14), *SPRED2* (6/14), *STK40* (6/14), and *INPPL1* (5/14). *INPPL1*, *MAP4K5*, or *LZTR1* deletion reactivated ERK signaling. INPPL1-mediated sensitization to SHP2i required its NPXY motif but not its lipid phosphatase domain. MAP4K5 acted upstream of MEK via a kinase-dependent target(s), whereas LZTR1 showed cell-dependent effects on RIT and RAS stability. *INPPLI*, *MAP4K5*, or *LZTR1* deletion also conferred SHP2i resistance in mice. Our results reveal multiple SHP2i resistance genes, emphasizing the need for detailed understanding of the resistance landscape to arrive at effective combinations.

## Introduction

The RAS/ERK cascade is a key signaling pathway downstream of receptor tyrosine kinases (RTKs), cytokine receptors, and integrins. Hyperactivation of this pathway, caused by gene amplifications, chromosomal abnormalities, or mutations, is among the most common oncogenic events in human cancer (1, 2). Multiple inhibitors targeting pathway components, including aberrant RTKs (e.g., EGFR, HER2, MET) or fusion-RTKs (e.g., BCR-ABL, FLT3-ITD), KRAS^G12C^ or BRAF^V600E^, and wild type MEK or ERK (3, 4), have been developed. These drugs can evoke dramatic tumor regressions in some patients and prolong their survival. Inevitably, however, resistance emerges, resulting in patient relapse and, in the absence of other, more durable modalities (e.g., immune therapies), patient demise (5–8).

Mechanisms of resistance to RAS/ERK cascade inhibitors are multiple and complex (5–8). Often, mutations in the target protein disable inhibitor binding. Alternatively, bypass mutations can occur in genes encoding downstream signaling components or components of parallel pathways. In addition, epigenetic alterations can induce a change in cell state (e.g., epithelial to mesenchymal transition, lineage switching) that renders malignant cells agnostic to the targeted oncogene. Some tumor cells treated with RAS/ERK pathway inhibitors exhibit “adaptive resistance,” in which ERK inhibition results in de-repression of MYC targets, including RTKs and their ligands, which drive increased upstream signaling and overcome inhibitor action (9–17).

The protein tyrosine phosphatase SHP2, encoded by *PTPN11*, is required for full activation of the RAS/ERK pathway (18–20). SHP2 functions upstream of RAS via as yet unknown substrate(s) to promote the activity of the guanine nucleotide exchange factors SOS1/2, although it might also have a role in RAS-GAP regulation and/or in parallel pathways. For these reasons, SHP2 inhibition has been proposed as a therapeutic strategy for malignancies driven by RTKs or fusion RTKs, “cycling” KRAS mutants, or, in combination with other pathway inhibitors for combating adaptive resistance (8, 11, 13, 16, 17, 21). Recently, allosteric inhibitors of SHP2 (SHP2i) were developed, and several have entered clinical trials as monotherapies or in various combinations (22–25). Because prospective identification of potential routes to drug resistance could suggest more effective combination therapies, we sought to identify potential mechanisms of SHP2i resistance using genome-wide loss-of-function screens.

## Results

### Genome-wide loss-of-function screens for SHP099 resistance

To systemically identify key regulators whose loss leads to SHP2i resistance, we performed genome-wide CRISPR knock-out screens on two FLT3-ITD-driven human AML lines, MOLM13 and MV4-11. These lines were chosen based on their marked sensitivity to SHP2 inhibition. We transduced each line with the TKOv3 lentiviral CRISPR library targeting 18,053 protein-coding genes with 4 guide RNAs per gene (26, 27) at 1,000X representation per replicate (n = 2 biological replicate screens per cell line). Cells were then cultured in vehicle (DMSO) or with the tool inhibitor SHP099 at 7X IC50 for 12 doublings (**Figure 1A**). The prolonged culture period and high concentration of inhibitor was designed to enrich for *bona fide* drivers of drug resistance, as opposed to synthetic lethal genes. At the end of the incubation period, genomic DNA (gDNA) was isolated from each sample and sequenced. Enrichment/dropout of individual genes was assessed by consistent effects on sgRNAs targeting the same gene using robust rank aggregation, as implemented in the MaGeCK (28) package (**Figure 1B-C, Supplemental Table 1**).

**Figure 1.**
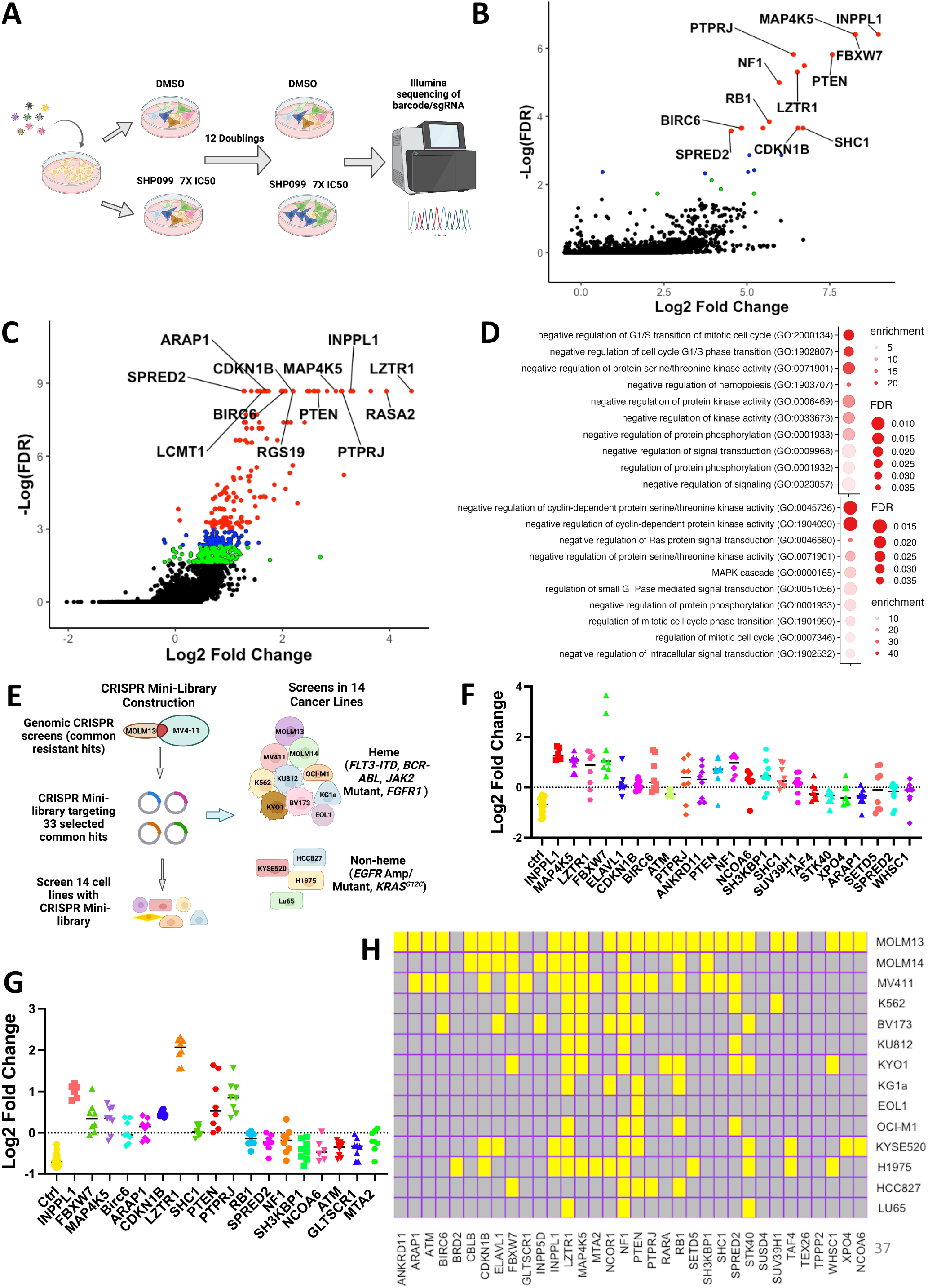
CRISPR screens identify resistant hits to SHP099. (A) Workflow for genome-wide CRISPR KO screens. **(B-C)** Results of CRISPR KO screen in MOLM13 (**B**) and MV4-11(**C**) cells, analyzed by MaGeCK; red circles indicate enriched genes with FDR<0.05, blue circles label enriched genes with 0.05<FDR<0.1, and green circles show genes with 0.1<FDR<0.2. **(D)** Gene Ontology analysis of top 50 resistance genes from genome wide MOLM13 (top panel) and MV4-11 (bottom panel) screens. Selected pathways (colored red in Supplemental Table 2) are shown. **(E)** Workflow for focused CRISPR mini-screens in 14 cancer lines. **(F-G)** Results of CRISPR mini-screens of MOLM13 (**F**) and MV4-11 (**G**) cells. Points represent log2-fold enrichment (SHP099 vs DMSO) of each sgRNA present in the library targeting the indicated gene, compared with non-targeting (ctrl) sgRNAs; significance was assessed by Student’s t test with FDR correction. **(H)** Heat map showing significant hits in CRISPR mini-screens across 14 cancer lines. Yellow color indicates FDR<0.05 enrichment for a given gene in a cell line.

Resistance genes that scored in both cell lines included known negative regulators of the RAS/ERK pathway (e.g., *NF1*, *SPRED2*, etc.), negative regulators of parallel pathways (e.g., *PTEN*), negative regulators of the cell cycle (e.g., *CDKN1B*, *FBXW7*, *RB1*, etc.), and *PTPRJ*, which encodes a receptor tyrosine phosphatase reported to target FLT3 (29, 30). We also found more unexpected genes, including *INPPL1*, *MAP4K5*, *LZTR1*, *BIRC6*, and the epigenetic regulators *SUV39H1*, *WHSC1*. (**Figure 1B-C, Supplemental Table 1**). Gene ontology (GO) analysis of the 50 top-ranked genes from MOLM13 and MV4-11 cells revealed enrichment for genes with annotations “negative regulation of protein serine/threonine kinase activity”, “negative regulation of protein phosphorylation” (MOLM13 and MV4-11), “negative regulation of cell cycle G1/S phase transition” (MOLM13), “negative regulation of cyclin-dependent protein kinase activity” (MV4-11), and “negative regulation of RAS protein signal transduction” (MV4-11), among others (**Figure 1D**, **Supplemental Table 2**). These results comport with known functions of the SHP2 pathway.

### Focused CRISPR screens identify genes that confer SHP099 resistance in multiple cancer cell lines

We next asked how frequently resistance genes from MOLM13 and MV4-11 cells were shared across a larger panel of cancer cells bearing different driver mutations. To this end, we constructed a focused CRISPR “mini-library” comprising lentiviruses expressing sgRNAs targeting 33 common hits from the above screens and several control sgRNAs (see Methods for details). Fourteen cancer cell lines were transduced with the mini-library in two independent experiments and cultured in vehicle or SHP099 (7X IC50) for 10 doublings (**Figure 1E**). Lines were chosen based on their *PTPN11* dependency in DepMap (31, 32), their observed sensitivity to SHP099 in our hands, and their diverse driver mutations (**Supplemental Table 3**). Most of these were AML (n=10, driven by FLT3-ITD, BCR-ABL, or JAK2^V617F^) lines; in addition, a few solid tumor lines driven by mutant EGFR or RAS^G12C^ were assessed (n=4). The distribution of sgRNAs was determined by sequencing of gDNA (**Supplemental Figure 1A**), and enrichment of each gene at the experimental endpoint was evaluated by one-tailed Student’s *t* test with FDR<0.05 considered significant. Replicates were well correlated (**Supplemental Figure 1B**), and non-targeting (control) guides were depleted as expected (**Figure 1F-G** and **Supplemental Figure 1C**). Several of the common resistance hits were either tumor suppressor genes or encoded negative regulators of the cell cycle (e.g., *PTEN, NF1, FBXW7*, *CDKN1B*, *RB1*). Germ line mutations in the negative regulators *NF1*, *SPRED2*, and *LZTR1* also cause RASopathies (33, 34). Others, such as *INPPL1* and *MAP4K5*, have no clear role in RAS/ERK pathway regulation or in cancer.

Across the 14 lines, the largest number of significant genes (FDR < 0.05) were found in MOLM13 and MV4-11 cells, which is to be expected given that the focused CRISPR library was designed based on our genome-wide screens of these cell lines (**Figure 1H**). Deletion of several genes conferred SHP099 resistance in multiple cell lines, including *LZTR1* (12/14), *NF1* (12/14), *PTEN* (8/14), *MAP4K5* (8/14), *STK40* (6/14), *FBXW7* (6/14), *SPRED2* (6/14), and *INPPL1* (5/14). Some hits were common to hematopoietic and solid tumor lines (e.g., *LZTR1, NF1, PTEN, STK40, INPPL1, MAP4K5*) and might predict frequent mechanisms of resistance in patients treated with SHP2is. The higher the SHP099 dose, the fewer significant hits that were recovered (**Supplemental Figure 1D**). Hence, confirmed hits from this screen (performed at 7X IC50) are likely to be genes whose absence causes substantial resistance to SHP099. We decided to further explore the mechanism by which the deletion of *LZTR1*, a near-universal hit, as well as two unexpected hits, *INPPL1* and *MAP45K*, confer SHP099 resistance.

### Structural basis for SHP2i resistance caused by *INPPL1* deletion

We first attempted for confirm that *INPPL1* deletion with either of two sgRNAs conferred SHP099 resistance *in vitro*. Parental and *INPPL1*-deleted (KO) MOLM13, MOLM14, and MV4-11 cells were cultured under standard conditions in the presence or absence of SHP099, and cell number was inferred by PrestoBlue assay. KO cells showed substantially higher readings, consistent with higher cell number. Importantly, re-expressing WT *INPPL1* in KO MOLM13 cells restored their sensitivity to SHP099, ruling out off target effects of the *INPPL1* sgRNAs as the cause of SHP099 resistance (**Figure 2A, right panel**). *INPPL1* KO also caused resistance to TNO-155 and RMC2550, two other SHP2 inhibitors, confirming that the effects of SHP099 on cell proliferation, and the resistance conferred by *INPPL1* deficiency, reflected SHP2 inhibition rather than off-target effects of the tool compound **(Supplemental Figure 2**). DAPI staining revealed that SHP099 causes G1 arrest and cell death (sub-G0 cells), both of which are decreased in KO cells (**Supplemental Figure 3**).

**Figure 2.**
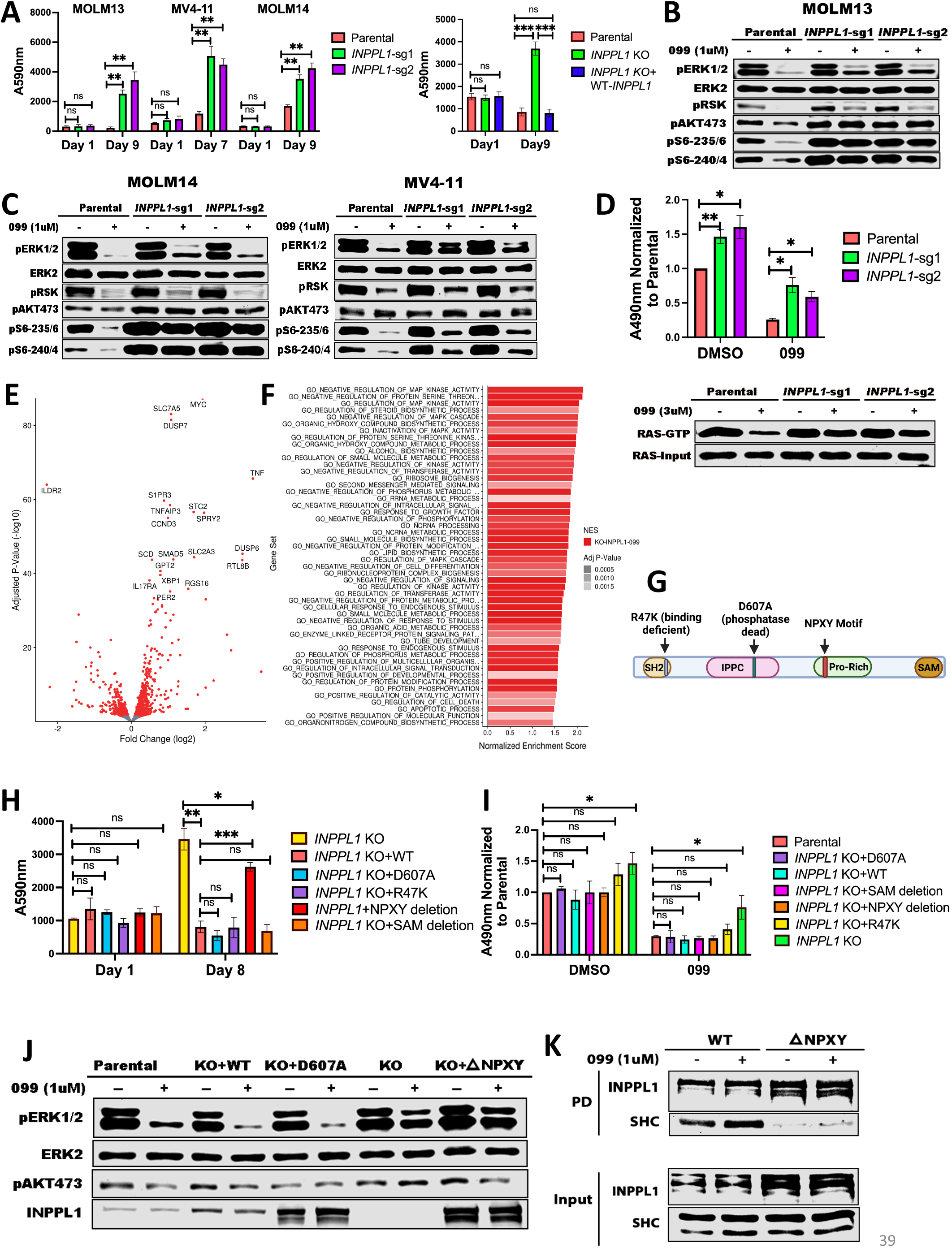
*INPPL1* KO acts downstream of RAS to promote ERK reactivation and SHP099 resistance. **(A)** *INPPL1* KO causes resistance to SHP099. Left panel shows results of PrestoBlue proliferation assays of MOLM13, MOLM14, and MV4-11 cells, treated with 1 µM, 2 µM, or 2 µM SHP099, respectively. Right panel shows proliferation of parental, *INPPL1* KO, and *INPPL1* KO re-expressing WT *INPPL1* MOLM13 cells in 1 µM SHP099. **(B-C)** Immunoblots of lysates from parental and *INPPL1* KO MOLM13 (**B**), MOLM14 (**C**), or MV411 (**C**) cells treated with vehicle or SHP099 (1 µM) for 1h; note increased activity of ERK pathway (pERK, pS6-235/6) in SHP099-treated *INPPL1* KO cells. **(D)** *INPPL1* KO causes increased RAS activation in MOLM13 cells. Cells were treated with vehicle (DMSO) or SHP099 (3 µM) for 1h, and RAS-GTP was quantified by active RAS ELISA (Cytoskeleton). Luminescence at A490 nm was normalized to parental DMSO values. Lower panel shows RAS RBD-pulldown assays on MOLM13 cell lysates. **(E)** Volcano plot showing significantly up- and down-regulated genes (q<0.05, red dots) in RNAseq analysis of *INPPL1* KO MOLM13 cells treated with SHP099 (1 µM) for 2.5 h compared with parental controls. **(F)** Top 50 significantly different MSigDB GO biological processes in *INPPL1* KO vs. parental MOLM13 cells, based on differentially expressed genes in RNAseq. **(G)** Domain structure of INPPL1 indicating mutants /deletions of functional domains assessed. **(H)** NPXY motif but not lipid phosphatase activity of INPPL1 is required to restore SHP099 sensitivity. WT *INPPL1* and the indicated *INPPL1* mutants were re-expressed in *INPPL1* KO MOLM13 cells, and cell number after exposure to SHP099 (1 µM) for 8 days was inferred by PrestoBlue assays. **(I)** NPXY motif is dispensable for INPPL1 effects on RAS activation. WT *INPPL1* and the indicated *INPPL1* mutants were re-expressed in *INPPL1* KO MOLM13 cells. Cells were treated with vehicle (DMSO) or SHP099 (3 µM) for 1h, and RAS activation was assessed by ELISA. All readings are normalized to vehicle-treated parental MOLM13 cells. **(J)** NPXY motif is required for ERK regulation by INPPL1. Parental or *INPPL1* KO MOLM13 cells reconstituted with the indicated mutants were treated with SHP099 (1 µM) for 1h, and cell lysates were prepared and analyzed by immunoblotting with the indicated antibodies. **(K)** *INPPL1* KO MOLM13 cells were reconstituted with WT- or NPXY deleted-INPPL1 bearing a C-terminal Strep tag. Cells were treated with vehicle (DMSO) or SHP099 (1 µM) for 1h, and INPPL1 was recovered on Strep-Tactin beads and analyzed by immunoblotting as indicated, *p<0.05, **p<0.01, ***p<0.001.

*INPPL1* encodes a protein also known as SHIP2, an SH2-domain containing 5’ phosphatidyinositol phosphatase expressed broadly in hematopoietic and non-hematopoietic tissues (35–38). SHIP2 catalyzes the dephosphorylation of PtdIns(4, 5)*P*_2_, PtdIns(3,4,5)*P*_3_, PtdIns(3, 5)*P*_2_, and several water soluble phosphoinositides, and is best known as a negative regulator of insulin-stimulated AKT activation. We explored how *INPPL1* deficiency causes SHP2i resistance. As expected, we observed increased levels of pAKT-S473 in *INPPL1*-KO MOLM13 cells in the presence or absence of SHP099. Activation of the AKT pathway results in increased mTORC1 activity, and ultimately increased activity of p70S6 kinase (S6K). Phosphorylation of S240/244 in ribosomal protein S6 (pS6-240/44), a selective target of S6K (39, 40), also was elevated in KO cells, providing additional evidence for AKT/mTORC1 pathway activation (**Figure 2B-C**). Unexpectedly, ERK, and its downstream target RSK showed increased phosphorylation in *INPPL1*-KO cells, as did S235/236 of S6 (pS6-235/6), which can be phosphorylated by S6K or RSK. Whereas SHP099 treatment strongly suppressed ERK, RSK, and S235/236 phosphorylation, suppression was attenuated in KO cells. SHP099 also resulted in decreased RAS activation (RAS-GTP) in parental cells, and this effect was attenuated by *INPPL1* KO. In addition, RAS activation was enhanced in vehicle-treated KO cells, indicating that *INPPL1* has an important role in negative regulation of RAS in these cells (**Figure 2D**).

To gain further insight into the predominant pathway(s) that confer SHP099 resistance, we performed RNAseq on parental and SHP099-treated MOLM13 cells (**Figure 2E** **and Supplemental Table 4**). In vehicle (DMSO)-treated cells, GO pathways most significantly enriched in *INPPL1* KO (compared with parental) cells featured annotations such as “biosynthetic process” and “metabolic process” (**Supplemental Figure 4A**), pathways that are regulated by the AKT/mTORC1 pathway (40). By contrast, in the presence of SHP099, *INPPL1*-KO cells predominantly showed enrichment for MAPK pathway gene sets (e.g., “Regulation of MAPK”, “Regulation of MAPK Cascade”, “MAPK Kinase Activity”). Moreover, “MYC genes”, “DUSP family”, and “SPRY family” genes are known targets of ERK, and were significantly upregulated in *INPPL1*-KO, compared with parental, cells (**Figure 2E-F**). Taken together, these findings suggest that although the AKT/mTORC1 and RAS/ERK pathways both show enhanced activation upon *INPPL1* deletion (and therefore are negatively regulated by INPPL1), increased activation of the former contributes predominantly to basal transcription (i.e., in the presence of vehicle), but increased RAS/ERK pathway activation likely is the major reason for SHP2i resistance.

INPPL1 is composed of several well-characterized domains (**Figure 2G**), including an N-terminal SH2 domain, a lipid phosphatase domain, a proline-rich region containing a NPXY motif that binds PTB-domain containing proteins, and a C-terminal SAM domain, which reportedly mediates protein-protein interactions (35–38). We asked which domain(s) is/are critical for INPPL1 to confer SHP2i sensitivity by restoring expression of WT *INPPL1* and various *INPPL1* mutants to *INPPL1* KO MOLM13 cells and assessing proliferation in the presence of SHP099 (**Figure 2H**). All constructs were overexpressed, but to comparable levels (**Figure 2J**). Only the NPXY motif deletion phenocopied the effects of *INPPL1* KO; remarkably, the ability of INPPL1 to confer sensitivity to SHP099 was independent of its lipid phosphatase activity (41), but instead appears to require its ability to bind (an)other protein(s) via its NPXY motif. Notably, expression of NPXY motif-deleted *INPPL1* restored activated RAS to levels comparable to WT *INPPL1*-expressing cells treated with SHP099, but ERK phosphorylation remained elevated (**Figure 2I-J**). These findings suggest that although INPPL1 regulates RAS in MOLM13 cells, its effects on SHP2 sensitivity are mediated downstream of RAS but upstream of ERK via an NPXY-binding protein(s). SHC binds to the NPXY motif (42–44), and, as expected, NPXY deletion greatly compromised SHC/INPPL1 co-immunoprecipitation (**Figure 2K**). Notably, SHC also scored as a resistance “hit” (FDR<0.05) in our genomic and mini-screens of MOLM13 and MV4-11 cells (**Figure 1B-C, F-H, Supplemental Table 1)**. These results suggest that a SHC/INPPL1 complex is required to confer SHP099 sensitivity in these cells, presumably by recruiting one or more additional proteins to limit ERK activation downstream of RAS.

### *MAP4K5* KO promotes resistance downstream of RAS via its kinase activity

*MAP4K5* (a.k.a. *KHS*, *GCKR*) encodes a member of the MAP4K family best known for its positive role in JNK activation (45, 46). Several MAP4K family members also can phosphorylate LATS1/2, resulting in Hippo pathway activation (47). However, the function of MAP4K5 is less clear. Using two independent sgRNAs, we first confirmed that *MAP4K5* deletion caused resistance to SHP099 (and other SHP2is) in MOLM13, MV4-11, KU812, KYO1 cells (**Figure 3A, Supplemental Figure 2**). Similar to the effects of *INPPL1* deletion, SHP099-treated *MAP4K5* KO cells showed reduced G1 arrest and decreased sub-G0 populations compared to parental controls (**Supplemental Figure 3**). The effect of *MAP4K5* deletion was unique among the *MAP4K*s, even though all members of the family were expressed in these lines (**Supplemental Figure 5A-B**). Re-expressing WT *MAP4K5,* but not a kinase-defective mutant (45), in KO MOLM13 cells restored SHP099 sensitivity (**Figure 3A-F** **and Supplemental Figure 5C**).

**Figure 3.**
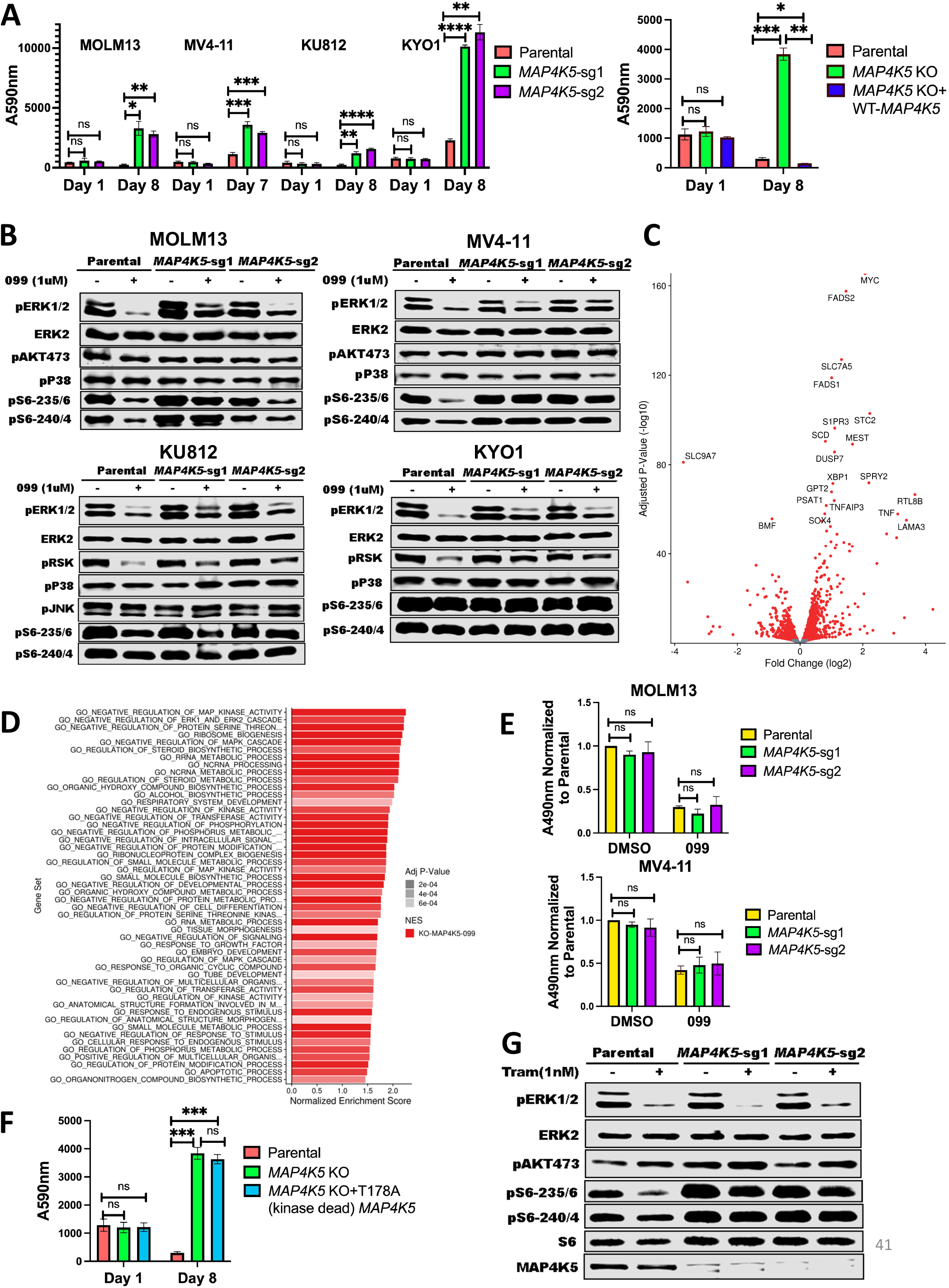
*MAP4K5* KO promotes SHP099 resistance downstream of RAS in a kinase-dependent manner. **(A)** *MAP4K5* KO causes SHP099 resistance. Left panel: PrestoBlue proliferation assays on MOLM13, MV4-11, KU812, or KYO1 cells treated with 1 µM, 2 µM, 5 µM, or 10 µM SHP099, respectively. Right panel: PrestoBlue assays of parental MOLM13 cells, *MAP4K5* KO MOLM13 cells, and *MAP4K5* KO MOLM13 cells reconstituted with WT *MAP4K5* in the presence of SHP099 (1 µM). **(B)** Increased ERK pathway activity in SHP099-treated MOLM13, MV4-11, KU812, and KYO1 cells lacking MAP4K5. Cells were treated with SHP099 (1 µM) for 1h, lysed, and analyzed by immunoblotting with the indicated antibodies. **(C)** Volcano plot showing significantly up- and down-regulated genes (q<0.05, red dots) in RNASeq analysis of *MAP4K5* KO MOLM13 cells treated with SHP099 (1 µM) for 2.5 h, compared with cognate parental controls. **(D)** Top 50 significantly different MSigDB GO biological processes in SHP099-treated *MAP4K5* KO vs. parental MOLM13 cells, based on differentially expressed genes from RNASeq. **(E)** *MAP4K5* KO does not affect RAS activation in MOLM13 and MV4-11 cells. MOLM13 and MV4-11 cells were treated with vehicle (DMSO) or SHP099 (3 µM MOLM13, 5 µM MV4-11) for 1h, lysed, and assessed for RAS activation by active RAS ELISA. **(F)** MAP4K5 requires its kinase activity to restore SHP099 sensitivity. Proliferation assays on parental MOLM13 cells, *MAP4K5* KO MOLM13 cells, or *MAP4K5* KO MOLM13 cells reconstituted with kinase dead MAP4K5 (T178A) in the presence of 1 µM SHP099. (**G**) MAP4K5 regulates ERK activation upstream of MEK. Parental or *MAP4K5* KO MOLM13 cells were treated with vehicle (DMSO) or the MEK inhibitor Trametinib (1 nM) for 1h, lysed, and analyzed by immunoblotting, *p<0.05, **p<0.01, ***p<0.001, ****p<0.0001.

In some cell lines tested, we were unable to detect activated JNK (pJNK) in the presence or absence of SHP099. However, even in a line in which pJNK was detectable (e.g., KU812), we observed no difference upon *MAP4K5* deletion, nor was p38 activation affected (**Figure 3B**). By contrast, SHP099-treated *MAP4K*5 KO cells bearing different RTK driver mutations, e.g., FLT3-ITD (MOLM13, MV4-11), and BCR-ABL (KU812, KYO1), showed increased pERK levels compared with cognate parental controls. Consistent with functional consequences of this increase in ERK activation, RNAseq revealed that the top enriched GO pathways in SHP099-treated *MAP4K5* KO cells (but not in vehicle-treated cells, **Supplemental Figure 4B**) had annotations including “Negative Regulation of MAP Kinase Activity” and “Negative Regulation of ERK1 and ERK2 Cascade”, and known ERK-responsive genes were upregulated (**Figure 3C-D** **and Supplemental Table 5**). In contrast to the effects of *INPPL1* deficiency, RAS activation was unaltered in vehicle- or SHP099-treated *MAP4K5* KO cells (**Figure 3E**). Taken together, these results indicate that MAP4K family members have redundant functions in JNK and p38 activation in MOLM13, MV4-11, KU812, and KYO1 cells, but MAP4K5 has a specific role in regulating ERK activation downstream of RAS. Consistent with this notion, *MAP4K5* KO cells are sensitive to MEKi treatment (**Figure 3G**).

### *LZTR1* KO promotes SHP099 resistance via cell-context dependent effects on RIT1 and/or RAS family members

Deletion of *LZTR1* using individual sgRNAs also caused resistance to SHP099 and other SHP2is in multiple cell lines (**Figure 4A**, **Supplemental Figures 2 and 3**). LZTR1 was reported by different groups to promote the degradation of RAS family members or RIT1 via recruitment of a CUL3 ubiquitin ligase complex (48, 49). Germ line *LZTR1* mutations also are responsible for some cases of the RASopathy Noonan syndrome (33, 34). Given the controversy over the target(s) of the LZTR1-containing ubiquitin ligase complex, we assessed the levels of RIT1 and RAS family members across multiple cell lines from our panel. Consistent with the observations of Castel *et al.* (49), RIT1 levels were increased in all *LZTR1* KO lines tested. Elevated MRAS levels were also seen in nearly all *LZTR1*-deficient lines (except KG1a). By contrast, increased levels of other RAS proteins were cell-dependent (**Figure 4B**). For example, in MV4-11 and KG1a cells, the levels of K-/N-/H-RAS were unaffected by *LZTR1* KO. By contrast, in K562 cells, LZTR1 deficiency led to increased levels of KRAS, HRAS, NRAS, and RRAS, in addition to the regularly seen elevation of RIT1 and MRAS (**Figure 4B**, **Supplemental Figure 6A**). K562 cells also showed increased levels of RAS-GTP (**Figure 4C**). The cognate transcripts for RAS family members were unaffected by LZTR1 deficiency, consistent with differential post-transcriptional regulation of RAS/RIT expression levels (**Supplemental Figure 6B**).

**Figure 4.**
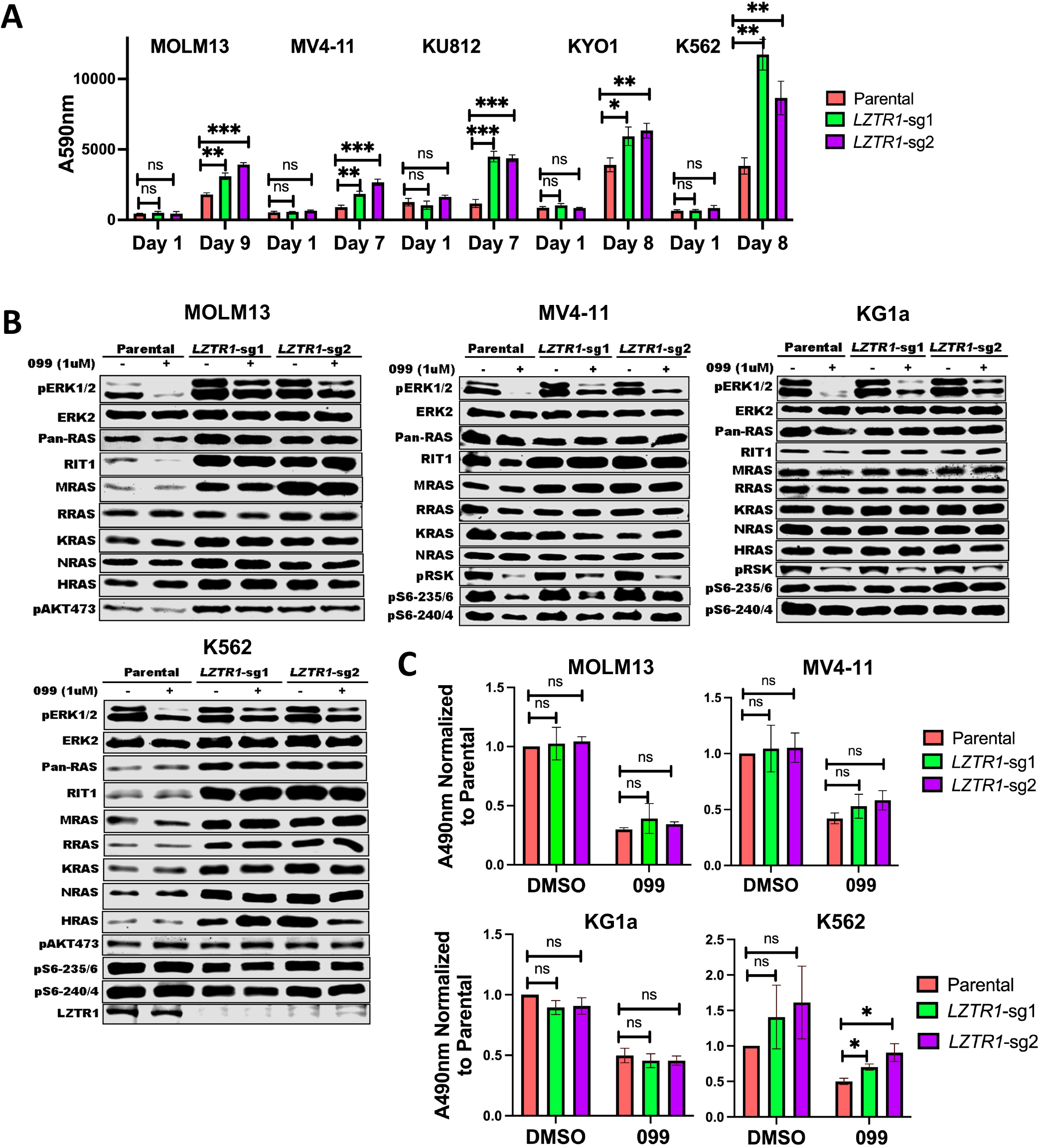
*LZTR1* KO promotes SHP099 resistance via cell-context dependent effects on RIT1 and/or RAS family members. **(A)** *LZTR1* KO cells are resistant to SHP099. PrestoBlue proliferation assays on parental or *LZTR1* KO MOLM13, MV4-11, KU812, KYO1, or K562 cells in 0.5 µM, 2 µM, 5 µM, 5 µM or 5 µM SHP099, respectively. **(B)** Cell context-dependent regulation of RIT and RAS family proteins by LZTR1. The indicated parental and *LZTR1* KO cells were treated with SHP099 (1 µM) for 1h, lysed, and analyzed by immunoblotting with the indicated antibodies. **(C)** *LZTR1* KO only increases active RAS levels in K562 cells. MOLM13, MV4-11, KG1a, and K562 cells were treated with vehicle (DMSO) or 3 µM (MOLM13) or 5 µM (other lines) SHP099 for 1h, and RAS-GTP levels were assessed by active RAS ELISA. All readings were normalized to those of vehicle-treated parental cells for each line, *p<0.05, **p<0.01, ***p<0.001.

These results indicate that RIT1 is a universal substrate for LZTR1 (followed by MRAS), whereas other RAS family members are conditional substrates based on the cell context (and potentially, differences in composition of LZTR1-containing ubiquitin ligase complexes; see Discussion). Regardless of whether RIT1/MRAS alone or additional combinations of RAS family members were increased, attenuation of ERK suppression upon SHP099 treatment was observed in all lines with *LZTR1* KO (**Figure 4B** and **Supplemental Figure 7B)**.

### *INPPL1*, *MAP4K5*, or *LZTR1* KO confer Gilteritinib resistance

Although SHP2 inhibitors are only in early phase trials, the FLT3 inhibitor Gilteritinib is used clinically for the treatment of FLT3-ITD^+^ AML (50, 51). MOLM13 and MV4-11 express FLT3-ITD, and our results indicate that INPPL1, MAP4K5, and LZTR1 act downstream of this fusion-RTK receptor. Therefore, we asked whether deletion of one or more of these SHP2i resistance hits also caused Gilteritinib resistance. Indeed, deletion of any of these genes caused resistance to Gilteritinib in both FLT3-ITD-expressing lines (**Figure 5A-B**).

**Figure 5.**
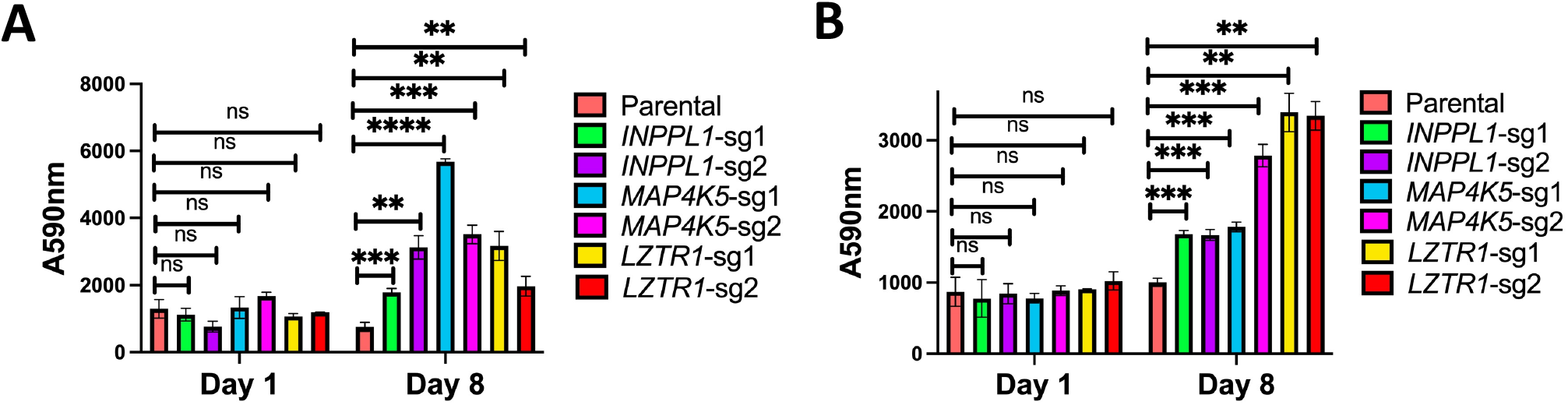
*INPPL1*, *MAP4K5*, or *LZTR1* KO also confer Gilteritinib resistance. (A) Parental MOLM13 cells and the indicated KO derivatives were treated with 2 nM Gilteritinib for 8 days and analyzed by PrestoBlue assays. **(B)** Same as **A**, but with parental and cognate KO MV4-11 cells with 0.6 nM Gilteritinib; **p<0.01, ***p<0.001, ****p<0.0001.

### *INPPL1*-, *MAP4K5*-, or *LZTR1*-KO confers SHP099 resistance *in vivo*

Finally, we tested whether the hits identified in our *in vitro* screen confer SHP099 resistance *in vivo*. To this end, MV4-11 cells were transduced with a lentivirus expressing luciferase to enable monitoring of tumor burden by bioluminescence imaging. We then generated *INPPL1-, MAP4K5-*, *or LZTR1-*deleted, luciferase-expressing lines by transduction with lenti-Cas9-expressing viruses co-expressing non-targeting or specific sgRNAs. MV4-11 cells and their derivatives (1.5 x 10^6^) were injected via tail veins into NSG mice. Ten days later, treatment with SHP099 (50 mg/kg/d) was initiated, and tumor burden was evaluated weekly by total body luminescence (**Figure 6A**). By week 3, mice bearing *INPPL1-, MAP4K5-, or LZTR1* KO MV4-11 cells all showed significantly higher tumor burdens than mice injected with parental cells (**Figure 6B-C**). Consistent with these data, *INPPL1*, *MAP4K5*, or *LZTR1* deletion also significantly shortened survival of injected mice (**Figure 6D**).

**Figure 6.**
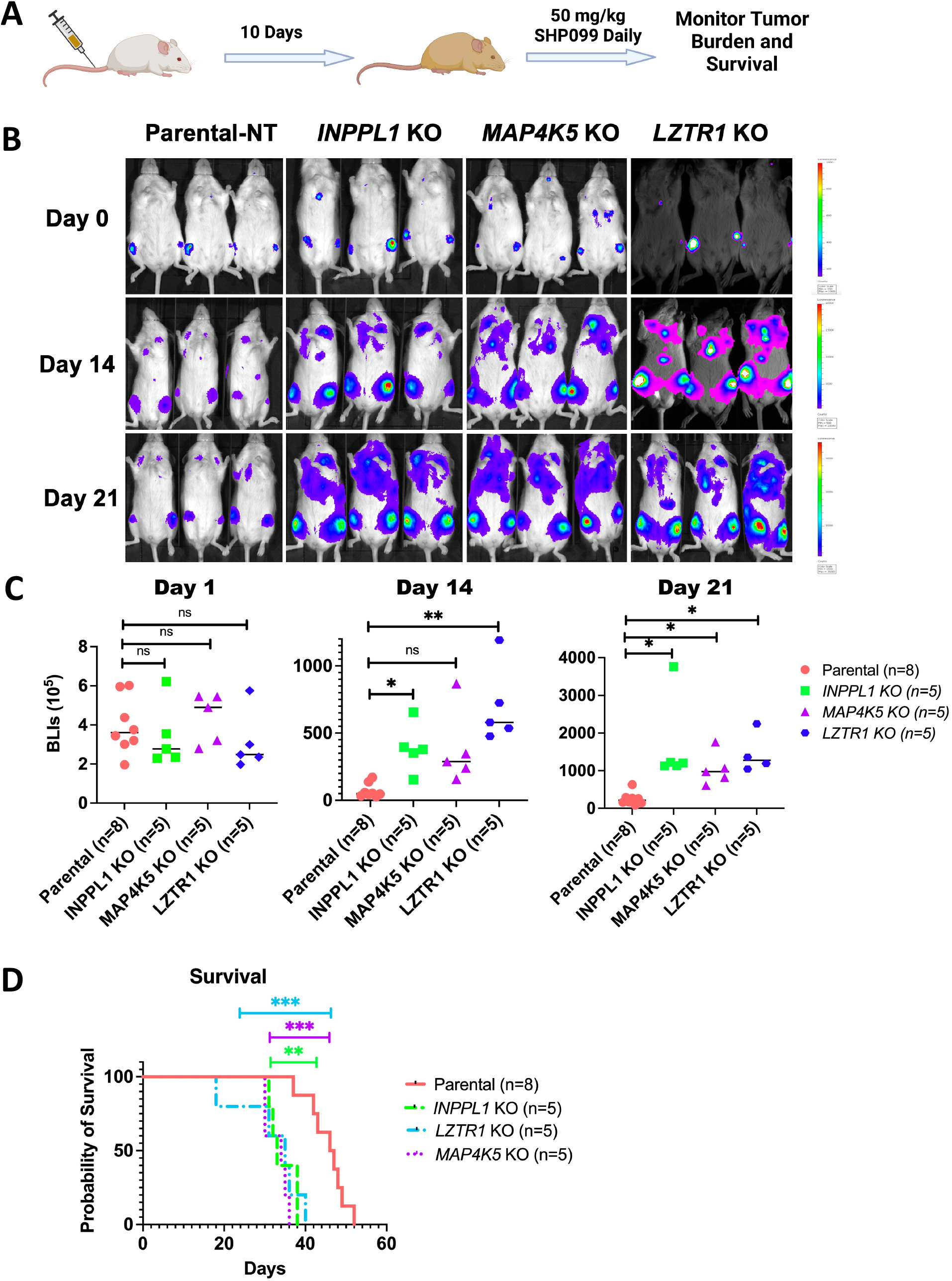
*INPPL1*-, *MAP4K5*-, or *LZTR1*-KO MV4-11 cells are resistant to SHP099 *in vivo.* **(A)** Workflow for mouse experiments. **(B)** Bioluminescence imaging of mice injected with parental and *INPPL1-*, *MAP4K5-*, *LZTR1*-KO MV4-11 cells on day 0 and monitored for 21 days. Note increased tumor burden caused by each KO line. **(C)** Quantification of data in **B** by Student’s t test. **(D)** Deletion of *INPPL1*, *MAP4K5,* or *LZTR1* in MV4-11 cells significantly shortens survival of mice, **p<0.01, ***p<0.001, Mantel log-rank test.

## Discussion

The essential role of SHP2 in RTK/RAS/ERK pathway signaling and the frequent activation of this pathway in multiple malignancies prompted the development of SHP2is for cancer therapy. Such inhibitors, alone or in combination with drugs targeting other cascade components or parallel pathways, are in clinical trials for RTK- or cycling KRAS mutant-dependent cancers. To anticipate SHP2i resistance mechanisms mimicking gene downregulation, silencing, or inactivation, we performed CRISPR/Cas9 genome-scale knock-out screens in two FLT3-ITD-driven human AML lines that are highly sensitive to SHP2 inhibition, followed by validation of common hits in 12 additional cancer lines. Our studies identified common and bespoke resistance mechanisms across these lines, including both “expected” and “novel” genes. The former included genes encoding known negative regulators of the RAS/ERK pathway (e.g., *NF1*, *SPRED2*), the parallel PI3K/AKT pathway (*PTEN*), and negative regulators of cell cycle (e.g., *CDKN1B*, *RB1*). Several of these genes were also found in previous genome-wide CRISPR loss-of-function screens for BRAF inhibitor resistance and metastasis (52, 53). However, more detailed analysis of three of the unexpected hits, *INPPL1* (SHIP2), *MAP4K5*, and *LZTR1*, reveal new insights into RAS/ERK pathway regulation. Deletion of each of these genes also results in SHP2i resistance in mice and cross-resistance to the FLT3-ITD inhibitor Gilteritinib.

INPPL1 is typically regarded as a negative regulator of the PI3K/AKT/mTORC1 pathway, but substantial evidence suggests additional, if not alternative, roles (28–31). Two *Inppl1* “knockout” mouse models exhibit increased activation of AKT and S6K, although the phenotypes of these mice differ, most likely reflecting the distinct parts of the *Inppl1* locus removed and/or removal of the adjacent *Phox2a* gene in one model (54, 55). Consistent with these findings, we observed increased pAKT and pS6-240/4 in *INPPL1*-KO cells. However, we also found that INPPL1 deficiency increased RAS/ERK activation, and RNAseq suggested that ERK pathway activation is important, if not dispositive, in driving SHP2i resistance. Notably, INPPL1 overexpression impairs ERK and/or AKT activation in response to multiple growth factors in various cell systems (44, 56–59). Decreased ERK activation in these experiments reflects sequestration of SHC from GRB2 by its binding to INPPL1 and thus might be an artifactual consequence of over-expression. More compellingly, zebrafish *Ship2* morphants show dorsal-ventral patterning defects characteristic of increased FGF signaling along with enhanced ERK phosphorylation and ERK-dependent gene expression (60). Moreover, the “pure” *Inppl1* knockout model (i.e., not including *Phox2a*) exhibits craniofacial abnormalities and runting, phenotypes that could reflect altered RAS/ERK activation (55). Likewise, a variety of homozygous inactivating mutations of human *INPPL1* cause opsimodysplasia, a rare skeletal dysplasia syndrome, rather than an insulin hypersensitivity syndrome (61–64). Taken together, it seems clear that INPPL1 can and does regulate the MEK/ERK pathway, at least in some cell contexts. Our results clearly reveal much more widespread, albeit non-ubiquitous, regulation of ERK activation by INPPL1.

The physiological function of INPPL1 lipid phosphatase activity also remains unresolved. Overexpression of catalytically inactive INPPL1 in smooth muscle cells suppresses growth factor-induced MEK/ERK activation, while enhancing AKT activation (58). As noted above, such results could reflect artifactual sequestration of SHC. By contrast, knock-in mice with global expression of catalytically impaired INPPL1 are runted and show craniofacial defects similar to those caused by global INPPL1 deficiency, as well as increased IGF1 (but not FGF2)-stimulated MEK/ERK activation. However, INPPL1 levels are significantly reduced in these mice, making it difficult to attribute the observed effects to catalytic impairment alone. Missense mutants affecting the phosphatase domain have been identified in Opsismodylpasia cohorts (61–64), but again, whether the mutant proteins are expressed at normal levels has not been established.

Our structure-function analyses show that phosphatase-inactive INPPL1 restores SHP099 sensitivity and decreases pAKT and pERK to the same extent as the WT protein. The SAM and Pro-rich domains are similarly dispensable. Instead, mutation of the NPXY motif, which enables binding to PTB domain-containing proteins, disables the ability of INPPL1 to restore SHP2i sensitivity to *INPPL1*-KO lines. SHC binds to this motif, and is the most likely mediator of the observed effects on ERK activation. Remarkably, however, the NPXY mutant restores the elevated RAS levels seen in KO cells to those of WT-reconstituted or parental cells, yet fails to normalize ERK phosphorylation. These results suggest that INPPL1 has a dual negative role in ERK regulation, acting upstream and downstream of RAS. Importantly, although the INPPL1 mutants were over-expressed compared with parental INPPL1 in our reconstitution experiments, expression of the phosphatase-dead D607A mutant (rescue-competent) and the NPXY deletion mutant (rescue-incompetent) were comparable.

MAP4K5 is one of six related STE-20-like kinases best known for their positive roles in JNK activation (45, 46). However, other functions have also been attributed to the MAP4K family. For example, MAP4K5 reportedly acts as a positive transducer of canonical and non-canonical WNT signaling in B cells (65). Other MAP4K family kinases can phosphorylate and activate LATS1/2 in HEK293 cells, acting in parallel to MST1/2 in the HIPPO pathway, although MAP4K5 was unable to perform this function (39). We find that MAP4K5, but not other MAP4K family members, negatively regulates the RAS/ERK pathway in a kinase-dependent manner, acting downstream of RAS but upstream of MEK; notably, we saw no difference in JNK activation in *MAP4K5*-KO cells. Although the mechanism by which MAP4K5 controls ERK activation remains unclear, recent biochemical studies found that MAP4K5 can phosphorylate all AMPK-related kinases (ARKs), including MARK1-3 (66). MARK1 (a.k.a. C-TAK1) phosphorylates KSR1 and impedes RAF/MEK/ERK activation (67). Studies are underway to test this interesting possibility.

Of the three resistance hits we characterized, *LZTR1* KO conferred SHP2i resistance the most widely (12/14 lines in the mini-screen). We did not directly test the remaining two lines individually, so we cannot exclude technical failure of *LZTR1* to validate in the mini-screen of these lines. Our results comport with previous studies demonstrating a role for LZTR1 in the RAS/MEK/ERK pathway, but also provide new insights into its mechanism of action. As an adaptor for CUL3 E3 ligase complexes, LZTR1 promotes the degradation of target proteins. However, the precise target (s) of the LZTR1/CUL3 complex has(ve) been a subject of controversy. RAS family members were suggested initially (48), but subsequent work argued that RIT1 is the primary substrate (49). We found that *LZTR1* KO invariably upregulated RIT1 and MRAS but not canonical RAS proteins in 10 cell lines screened. Nevertheless, in other cell lines, various RAS proteins also were upregulated, with K562 cells showing stabilization of KRAS, HRAS, NRAS, and RRAS. The simplest explanation for these findings is that LZTR1/CUL3 complexes have additional components that await identification. For example, an inhibitor of RAS protein binding could be present or a RAS-specific adaptor could be missing in lines where only RIT1 is stabilized. A very recent study of leukemogenesis in *Lztr1^-/-^* mice also noted altered levels of conventional RAS proteins along with RIT1 and MRAS (68).

Although we characterized three targets in detail in this work, several other novel hits warrant further study. *STK40*, which encodes a poorly understood pseudokinase, was a frequent hit (6/14 lines), whose deletion causes SHP099 resistance to in hematologic (*FLT3-ITD* and *BCR-ABL*-driven) and solid tumor (*EGFR*-driven) cells. We also recovered genes encoding epigenetic regulators, including *WHSC1*, *SETD5*, and *SUV39H1*, as well as transcriptional coactivators/repressors (*NCOR1*, *NCOA6*). While our screens were designed with the primary goal of identifying mechanisms of SHP099 resistance, we did observe some synthetic lethal genes. For example, *KDM5A,* which encodes a Jumanji-type histone demethylase, was depleted significantly in SHP099-treated MV4-11 and MOLM13 cells, and several additional genes showed synthetic lethal behavior in MV4-11 cells in particular.

Because SHP2is are in early-stage trials, clinical samples of resistance are not available. Nevertheless, several lines of evidence indicate that genes identified here are relevant to therapeutic outcome in hematologic and other malignancies. Deletion of *INPPL1*, *MAP4K5*, or *LZTR1* confers Gilteritinib resistance (**Figure 5A-B**); similar effects of *LZTR1* deficiency were reported recently (68). High levels of *MAP4K5* RNA are associated with favorable prognosis in the TGCA AML cohort, both in univariate analyses and when corrected for age, cytogenetics, FAB stage, and *RUNX* and *TP53* status (69). Conversely, low *MAP4K5* levels correlate with particularly poor outcome in pancreatic ductal adenocarcinoma (70). Future studies will be required to assess the generality of those relationships and their utility for combination therapy with SHP2is, as well as to determine whether deletion or mutation of the resistance hits observed in these studies are found in patients treated with SHP2is and other targeted therapies.

## Methods

### Cell lines and reagents

Cells were maintained in 5% CO2 at 37^0^C in media conditions described by the vendor or the source laboratory and were tested monthly for mycoplasma by PCR (71). KO cells were generated by infection with lentiviruses constitutively expressing Cas9 and appropriate sgRNAs. All experiments were performed with early passage lines within three months of defrosting. MOLM13, MV4-11, BV173, K562, H1975, LU65, HCC827 cells were from lab stocks. KYSE520 was obtained from the DSMZ (German Collection of Microorganisms and Cell Cultures GmbH). KG1a was obtained from Dr. Christopher Park (NYU Grossman School of Medicine) in June 2020; MOLM14 was obtained from Dr. Iannis Aifantis (NYU Grossman School of Medicine) in July 2020; KU812, KYO1, EOL1, OCI-M1 were provided from Dr. Ross L. Levine (Memorial Sloan Kettering Cancer Center) in March 2021. MOLM13, MV4-11, BV173, K562, H1975, LU65, HCC827, KYSE520, KG1a, KU812, KYO1 and EOL1 were cultured in RPMI supplemented with 10% FBS and 1% penicillin/streptomycin (P/S). OCI-M1 was maintained in IMDM supplemented with 10% FBS and 1% P/S. sgRNAs for non-targeting control and the target genes *INPPL1*, *MAP4K5* and *LZTR1* were as follows (target sequence):

Non-Targeting: AACCGGCTGCGCGTTTGCAA

*INPPL1*-sg1: GCAGGGCGCACACAAGGCCC

*INPPL1*-sg2: CCTGGATATCCATGTCCAGG

*MAP4K5*-sg1: AGGACTACGAACTCGTCCAG

*MAP4K5*-sg2: TAGGCCAGAAATGTACACAC

*LZTR1*-sg1: TATGGTCGAAGTCCACGCTC

*LZTR1*-sg2: CGGCCGAGTGGTGGTAACGG

Parental and KO MV4-11 cells for mouse experiments were generated with non-targeting, *INPPL1*-sg1, *MAP4K5*-sg1, and *LZTR1*-sg1. sgRNA resistant constructs for *INPPL1* and *MAP4K5* for re-expression were generated through making silent mutations in PAM sequence for *INPPL1*-sg1 and *MAP4K5*-sg1.

### Plasmids and lentiviruses

The open reading frames (ORFs) of *INPPL1* (GenScript: OHu19348), *MAP4K5* (GenScript: OHu02673), and various mutants of these genes were cloned into pLV-EF1a-IRES-Neo (Addgene: 85139) by using a Gibson Assembly kit (NEB: E5510S). All sgRNAs were cloned into lentiCRISPRv2 (Addgene: 52961) by following a published protocol (52, 72). Lentiviruses were produced by co-transfecting HEK293T cells with viral construct and packaging vector DNAs (26, 27). Cells were spin-infected with concentrated viruses and infected cells were recovered by selection with the appropriate antibiotics.

### Cell assays

For proliferation and dose-response assays, cell number was inferred by PrestoBlue assay (Thermo Fisher: A13262). Briefly, cells were seeded in 96-well plates and then incubated with media containing DMSO/various inhibitors. For proliferation assays, viability was assessed, and media (including inhibitors) were refreshed, every two days. For dose-response studies, media (including inhibitors) were refreshed once at Day 2, and data were collected at Day 4. Dose response curves (and IC50s) were generated with R package “drc”. To assess cell cycle distribution, cells were fixed with 70% ethanol overnight at -20^0^C, followed by DAPI staining the next day, and analyzed by flow cytometry using FlowJo software.

### CRISPR mini-library construction

A CRISPR mini-library was constructed based on overlapping hits from the genome wide screens of MV4-11 and MOLM13 cells. The library comprised 142 sgRNAs with 4sgRNA/gene against 33 target genes and 10 non-targeting controls. For the target genes, two sgRNAs were derived from the TKOv3 library and the other two were from the Doench CRISPR KO library and had the highest “Rule Set 2 Scores” (73). Mini-library sgRNAs were cloned into the lentiCRISPRv2 backbone according to previously published protocol (53) and sequenced to assess library quality (**Supplementary Figure 1A**).

### CRISPR/Cas9 screens

Two independent replicas of each cell line were spin-infected with TKOv3 Cas9-sgRNA or CRISPR mini-library viruses at low MOI (∼0.2) and at 1000X representation for each sgRNA in the library. At Day 2 post-infection, puromycin was added to the media, and cells were selected for 8 days. Live cells (for hematopoietic cell lines) after selection were isolated using Ficoll-Paque PLUS Media according to the manufacturer’s instructions, and cultured without antibiotics for additional 2 days. After recovery, a 1000X library representation of cells for each replica was cultured with DMSO or SHP099 at 7x IC50 for each line for 12 doublings (genomic screens) or 10 doublings (mini-screens). For mini-screens, if the IC50 for the cell line was >1.5 μM, 10 μM SHP099 was used. At screen termination, gDNA was extracted and PCR amplified according to published protocol (20–21). Final PCR products were sequenced with Illumina NovaSeq 6000 (SP 100 Cycle Flow Cell v1.5). Results were decomplexed with Bowtie (62) to generate count tables and subsequently analyzed with MaGeCK for genomic screen or t test with FDR correction for mini-screens.

### Genomic DNA extraction

Briefly, cells (3-5X10^7^) were suspended in 6 ml lysis buffer (50mM Tris, 50mM EDTA, 1% SDS, pH 8), supplemented with 30 µl 20mg/ml Proteinase K (Qiagen 19131), and incubated at 55°C overnight. For higher cell numbers, volumes were adjusted appropriately. The next day, 30 µl of 10mg/ml RNAse A (Qiagen 19101) were added to the sample, which was then incubated at 37°C for 30 mins. Then, proteins in the sample were precipitated with 2 ml 7.5M cold ammonium acetate and centrifuged at >4,000g for 10 mins. Supernatants containing DNA were transferred to a new tube, isopropanol (6 ml) was added, and the samples were centrifuged at >4000g for 10 mins. Precipitated DNA was washed with 6 ml 70% ethanol, air dried at room temperature, and suspended in nuclease-free water.

### Biochemical assays

RAS activation was assessed by ELISA using a commercially available kit (Cytoskeleton: BK131) according to manufacturer’s instructions, or by GST-RBD “pull-down” assays. For active RAS pulldown assays, MOLM13 cells were lysed in RAS PD buffer (50mM Tris-HCl, pH 7.4, 150mM NaCl, 5mM MgCl2, 5% glycerol, 1% NP40) supplemented with protease and phosphatase inhibitors on ice and centrifuged immediately at 14,000 rpm for 10 min at 4°C. Clarified lysates were incubated with GST-RBD glutathione agarose beads for 45 mins with constant rocking. Beads were washed 3 times with PD buffer and suspended in 2X SDS-PAGE sample buffer.

For monitoring additional effects on cell signaling pathways, 2X10^6^ cells/well were plated in 6-well plates, and the next day the indicated inhibitor(s) were added for 1h. Cells were harvested and lysed in RIPA buffer (50 mM Tris-HCl pH 8, 150mM NaCl, 2mM EDTA, 1% NP-40 and 0.1% SDS), supplemented with protease (40 µg/ml PMSF, 2 µg/ml antipain, 2 µg/ml pepstatin A, 20 µg/ml leupeptin, and 20µg/ml aprotinin) and phosphatase (10 mM NaF, 1 mM Na3VO4, 10 mM β-glycerophosphate, and 10 mM sodium pyrophosphate) inhibitors. Protein concentration was determined by Coomassie assay according to the manufacturer’s instructions (Abcam: #ab119211). For immunoblotting, total lysate protein (15 μg) was resolved by SDS-PAGE and transferred to nylon membranes in 1X transfer buffer supplemented with 10% methanol. Membranes were incubated with primary antibody (in 5% BSA in TBST) at 4^0^C overnight, and then with secondary antibodies labeled with IRDye (in 3% skim milk in TBS with 0.1% SDS/0.5% Triton) at room temperature for 1h, followed by visualization of bands with a LICOR Odyssey CLx apparatus.

Antibodies against p-p42/44 MAPK (#9101; 1:1000), p-AKT (Ser473) (#9271; 1:1000), p-S6 (Ser240/244) (#5364; 1:1000), p-S6 (Ser235/236) (#2211; 1:1000), total S6 (#2317; 1:1000), p-SAPK/JNK (Thr183/Tyr185) (#4668; 1:1000), p-p38 (Thr180/Tyr182) (#4511), p-RSK (359/363) (#9344, 1:1000), SHIP2 (#2839, 1:1000), and RRAS (#8446, 1:1000) were obtained from Cell Signaling. Antibodies against LZTR1 (sc-390166 X, 1:1000) and ERK2 (sc-1647; 1:1000) were obtained from Santa Cruz Biotechnology. Antibodies against MAP4K5 (#ab96551, 1:1000) and MRAS (#ab176570, 1:1000) were from Abcam. Pan-RAS antibody (Ab-3, 1:1000) was from Millipore. Antibodies against KRAS (12063-1-AP, 1:5000), NRAS (10724-1-AP, 1:1000), and HRAS (18295-1-AP, 1:1000), were from Proteintech.

A short peptide sequence (Strep-tag: ASWSHPQFEK) was cloned to C-terminus of WT and NPXY-deleted *INPPL1* and re-expressed in endogenous *INPPL1* KO MOLM13 cells for affinity purification. For pulldown of strep-tagged INPPL1, MOLM13 cells were lysed in NP40 buffer (50mM Tris-HCl, pH 7.4, 150mM NaCl, 1%NP40, 5mM EDTA). After clarification, lysates were incubated with MagStrep “type3” XT beads (#2-4090-010, IBA) overnight at 4 °C. The next day, beads were washed for 3 times with NP40 buffer, then suspended with 2X SDS-PAGE sample buffer, boiled at 95 °C for 10 mins, and analyzed by immunoblotting.

### RNA extraction and sequencing

For RNAseq sample preparation, parental, *INPPL1* KO, and *MAP4K5* KO MOLM13 cells (2 million cells/well) were plated in 6 well plates. The next day, cells were treated with DMSO or 1 μM SHP099 for 2.5h, and RNA was extracted immediately afterwards with a Qiagen RNeasy Plus Mini Kit (Qiagen: 74136).

RNA levels were quantified using RNA Nano Chips (Agilent: #5067-1511) on an Agilent 2100 BioAnalyzer. RNA-Seq library preps were constructed with the Illumina TruSeq® Stranded mRNA Library Prep kit (Illumina: #20020595) using 1000 ng of total RNA as input, amplified by 10 cycles of PCR, and sequenced paired-end 50 cycles on an Illumina NovaSeq6000 SP flowcell with a 2% PhiX spike-in.

Sequencing results were demultiplexed and converted to FASTQ format using Illumina bcl2fastq software. The sequencing reads were adapter- and quality-trimmed with Trimmomatic and then aligned to the human genome (build hg38/GRCh38) using the splice-aware STAR aligner. The featureCounts (74) program was utilized to generate counts for each gene based on how many aligned reads overlap its exons. These counts were then normalized and used to test for differential expression using negative binomial generalized linear models implemented by the DESeq2 R package. Pathway enrichment analysis was performed for the pre-ranked gene lists based on the differential expression using the fgsea R package and MSigDB gene sets. All sequence data are available in GEO (Accession number pending).

### qPCR methods

Total RNA was isolated by using the Qiagen RNeasy kit (Qiagen: 74136). cDNAs were generated by using the SuperScript IV First Strand Synthesis System (Invitrogen: 18091050). qRT-PCR was performed with Fast SYBR™ Green Master Mix (FisherScientific: 4385618), following the manufacturer’s protocol, in 96-well format in C1000 Touch Thermal Cycler (Biorad). Differential gene expression analysis was performed with CFX Manager (Biorad) and normalized to *GAPDH* expression. Primers used are listed in Supplemental Table 6.

### Mouse experiments

To assess the effects of resistance gene depletion on leukemogenesis, Mv4-11 cells were transduced with a luciferase-expressing lentivirus, and then infected with lentiviruses co-expressing Cas9 and sgRNAs for *INPPL1*, *MAP4K5*, *LZTR1* or a non-targeting control and selected with puromycin. Each cell line (1.5 million cells) was injected to NSG mice via tail vein, and 10 days later, SHP099 (50 mg/kg) via oral gavage daily was initiated. Whole-body bioluminescence imaging using an IVIS imager was performed at the indicated times immediately after retro-orbital injection of 150 mg/kg D-luciferin Firefly (Perkin-Elmer, #122799). Bioluminescence signals were quantified using Living Imaging software with standard regions of interest (ROI) rectangles.

### Statistical analysis

Results are expressed as mean ± SD, as indicated. Statistical significance was assessed by Student’s *t* test, for proliferation assays and active RAS ELISAs. Survival curves were plotted using the Kaplan-Meier algorithm and Prism software, and significance was assessed using the Log-rank (Mantel-Cox) test. A p<0.05 was considered statistically significant.

### Animal study approval

All the animal experiments were conducted in accordance with the Guide for the Care and Use of Laboratory Animals and approved by the Institutional Animal Care and Use Committees at New York University Grossman School of Medicine.

### Authors’ contributions

W. Wei: experimental design, experiments execution, and writing of the manuscript. M.J. Geer: generating data for *in vitro* and *in vivo* experiments, data analysis and editing of the manuscript. X. Guo: design, construction and evaluation of the CRISPR mini-library. I. Dolgalev: RNAseq data analysis and writing of the manuscript; N.E. Sanjana: advice in genome-wide CRISPR screens, construction of the mini-library; B.G. Neel: overall supervision of the project, data analysis, writing of the manuscript.

## Acknowledgements

We thank Drs. Julia Ban and Xufeng Chen for help with the mouse leukemogenesis experiments and Drs. Hao Ran and Carmine Fedele for technical advice and helpful discussions. We also thank the PCC Genome Technology Center, Applied Bioinformatics Laboratory, and Immune Monitoring Laboratory shared resources, funded by P30 CA016087. This work was supported by grants CA49152 and CA244896 to B.G.N. and HG010099 to N.E.S.

