## Supplementary for "Genome-wide CRISPR/Cas9 Screens Reveal Shared and Bespoke Mechanisms of Resistance to SHP2 inhibition"

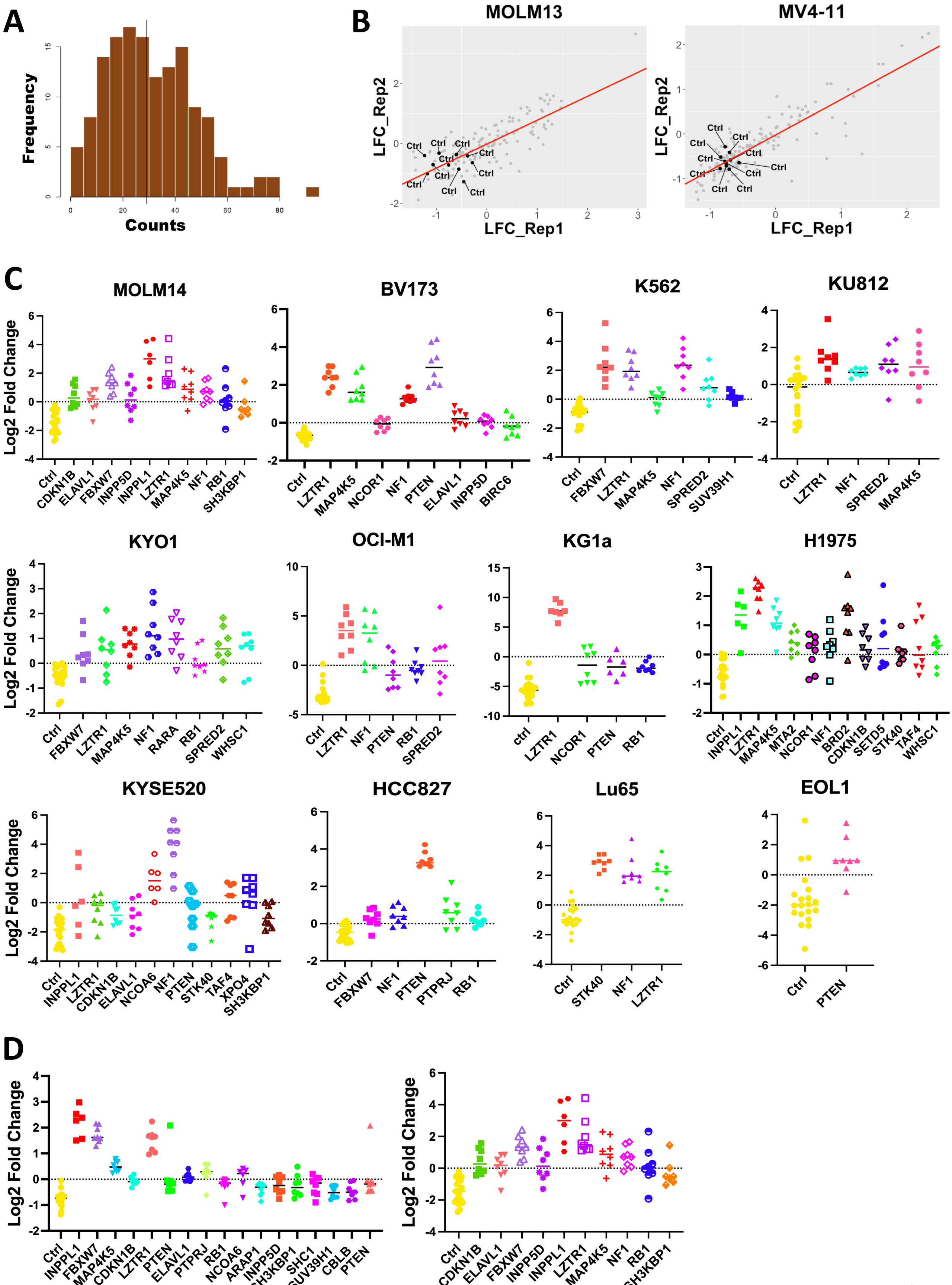
A
Frequency
Counts
B
MOLM13
MV4-11
LFC_Rep1
LFC_Rep2
Ctrl
C
MOLM14
Ctrl CDKN1B ELAVL1 FBXW7 INPP5D INPPL1 LZTR1 MAP4K5 NF1 RB1 SH3KBP1
BV173
Ctrl LZTR1 MAP4K5 NCOR1 NF1 PTEN ELAVL1 INPP5D BIRC6
K562
Ctrl FBXW7 LZTR1 MAP4K5 NF1 SPRED2 SUV39H1
KU812
Ctrl LZTR1 NF1 SPRED2 MAP4K5
KYO1
Ctrl FBXW7 LZTR1 MAP4K5 NF1 RARA RB1 SPRED2 WHSC1
OCI-M1
Ctrl LZTR1 NF1 PTEN RB1 SPRED2
KG1a
ctrl LZTR1 NCOR1 PTEN RB1
H1975
Ctrl INPPL1 LZTR1 MAP4K5 MTA2 NCOR1 NF1 BRD2 CDKN1B SETD5 STK40 TAF4 WHSC1
KYSE520
Ctrl INPPL1 LZTR1 CDKN1B ELAVL1 NCOA6 NF1 PTEN STK40 TAF4 XPO4 SH3KBP1
HCC827
Ctrl FBXW7 NF1 PTEN PTPRJ RB1
Lu65
Ctrl STK40 NF1 LZTR1
EOL1
Ctrl PTEN
Log2 Fold Change
D
Ctrl INPPL1 FBXW7 MAP4K5 CDKN1B LZTR1 PTEN ELAVL1 PTPRJ RB1 NCOA6 ARAP1 INPP5D SH3KBP1 SHC1 SUV39H1 CBLB PTEN
Ctrl CDKN1B ELAVL1 FBXW7 INPP5D INPPL1 LZTR1 MAP4K5 NF1 RB1 SH3KBP1
Log2 Fold Change

**Supplemental Figure 1. Focused CRISPR mini-screens identify genes conferring SHP099 resistance in 12 additional cell lines. (A)** sgRNA distribution in cloned CRISPR mini-library: 141/142 sgRNAs were detected. **(B)** Excellent correlation between replicate CRISPR mini-screens of MOLM13 and MV4-11 cells. The Log2 fold enrichment of each sgRNA is shown for treatment (SHP099 at 7X IC50 for each line) vs vehicle (DMSO) groups at screen termination. All non-targeting sgRNAs (Ctrl) were depleted as expected. **(C)** Significantly enriched genes (FDR<0.05) in CRISPR mini-screens of the indicated cancer cell lines. **(D)** Effect of SHP099 concentration on recovery of resistance genes in MOLM14 cells. Left panel: 5X IC50 (3 μM), Right panel: 7XIC50 (5 μM).

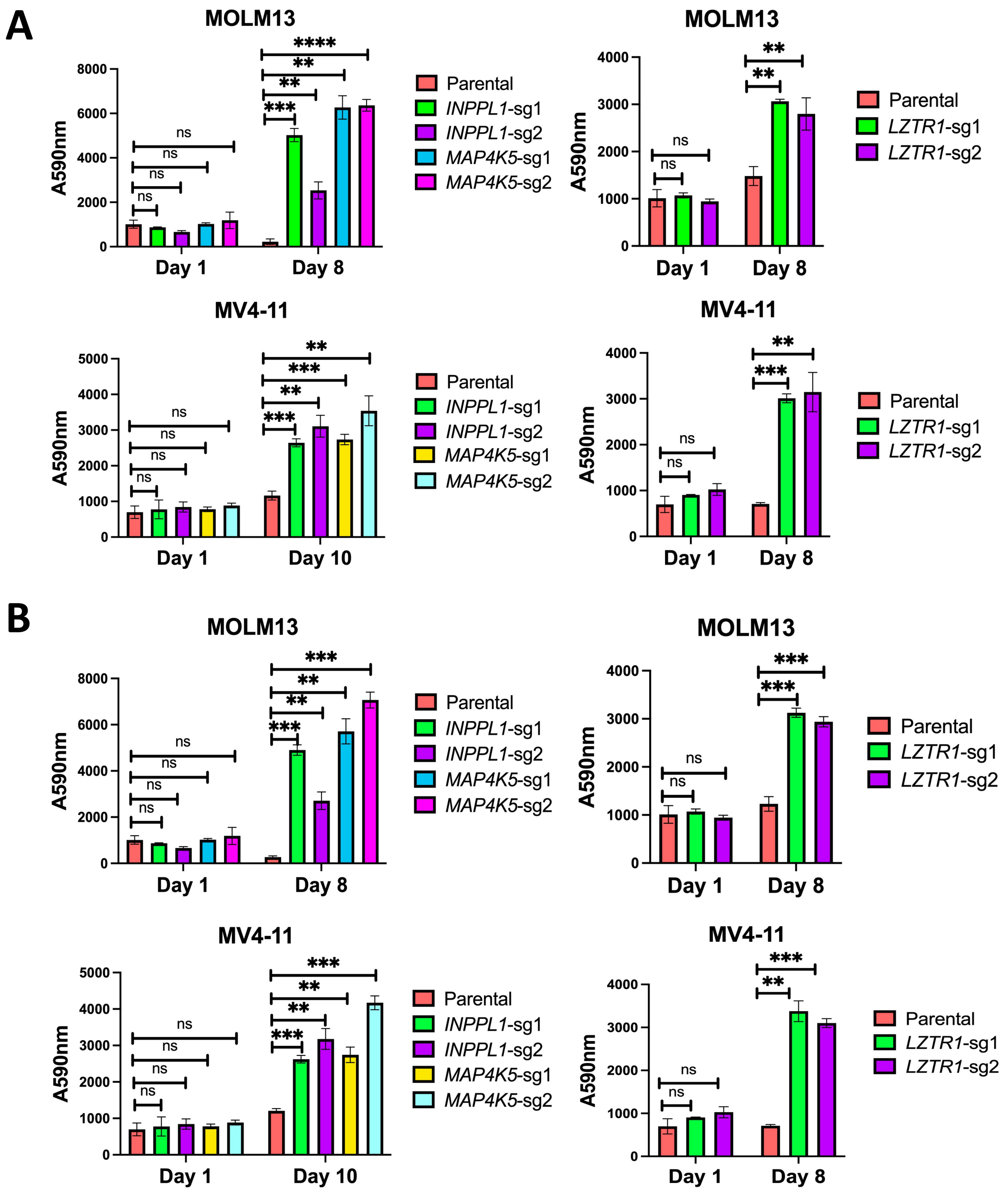


**Supplemental Figure 2. *INPPL1*, *MAP4K5*, or *LZTR1* KO confer resistance to other SHP2 inhibitors. (A)** Top left panel: PrestoBlue proliferation assays on parental, *INPPL1* KO, or *MAP4K5* KO MOLM13 cells treated with RMC-4550 (60 nM); Top right panel: PrestoBlue proliferation assays on parental or *LZTR1* KO cells treated with RMC-4550 (30 nM). Bottom left panel: PrestoBlue proliferation assays on parental, *INPPL1* KO, or *MAP4K5* KO MV4-11 cells treated with RMC-4550 (200 nM); Bottom right panel: PrestoBlue proliferation assays on parental or *LZTR1* KO MV4-11 cells treated with RMC-4550 (300nM). **(B)** Same design as in **A**, but with TNO155. Doses: 60 nM for parental and *INPPL1* KO/*MAP4K5* KO MOLM13 cells; 30 nM for parental and *LZTR1* KO MOLM13 cells, or 200 nM for parental and *INPPL1* KO/MAP4K5 KO MV4-11 cells or 300 nM for parental and *LZTR1* KO cells. **p<0.01, ***p<0.001, ****p<0.0001.

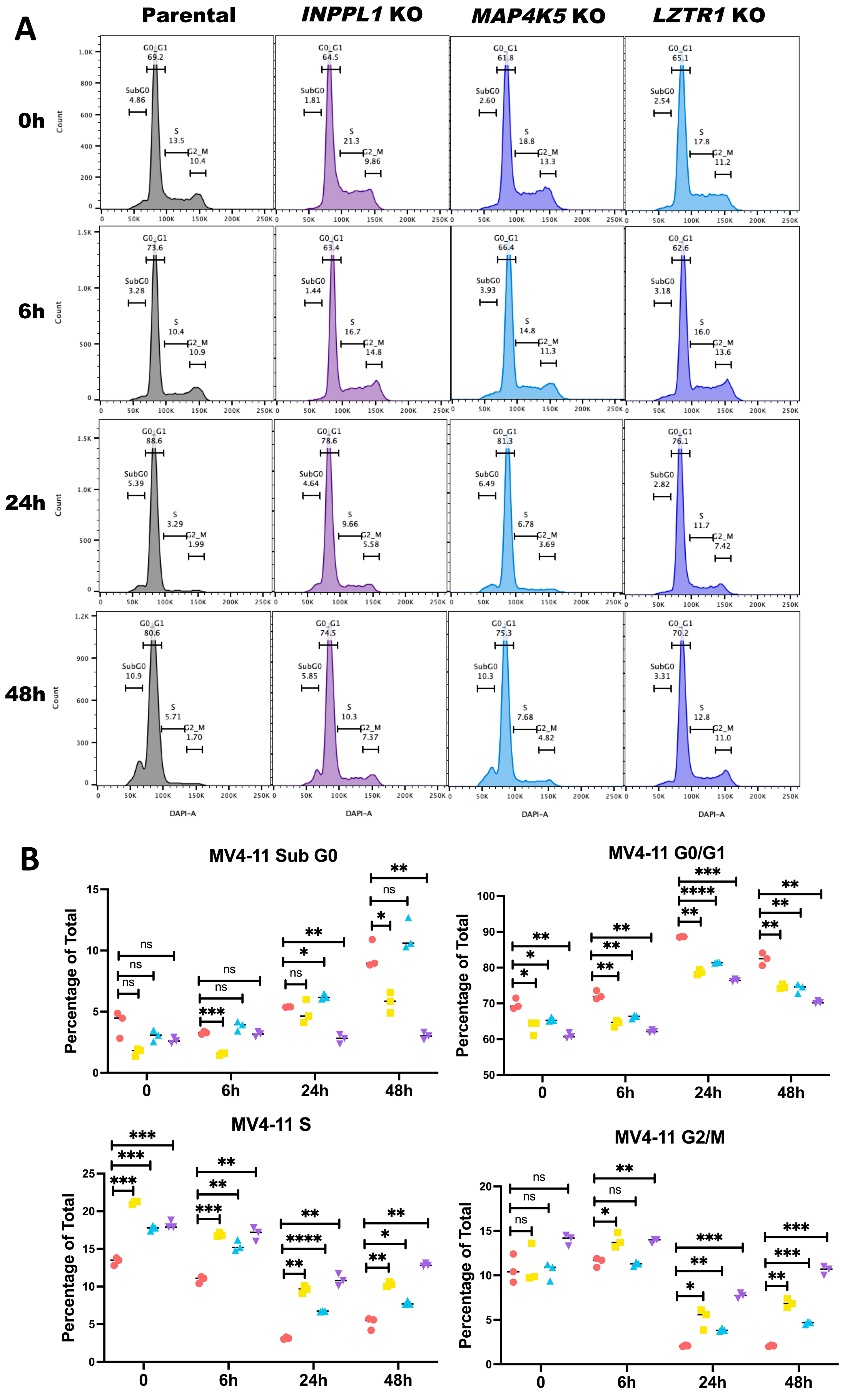
A
Parental
INPPL1 KO
MAP4K5 KO
LZTR1 KO
0h
6h
24h
48h
Count
DAPI-A
SubG0
G0_G1
S
G2_M
B
MV4-11 Sub G0
MV4-11 G0/G1
MV4-11 S
MV4-11 G2/M
Percentage of Total
0
6h
24h
48h
Parental
INPPL1 KO
MAP4K5 KO
LZTR1 KO

**Supplemental Figure 3. Effects of *INPPL1*, *MAP4K5*, or *LZTR1* deletion on cell cycle response to SHP099 in MV4-11 cells.** (**A)** Cell cycle distribution of parental, *INPPL1*, *MAP4K5* and *LZTR1* KO MV4-11 cells treated with vehicle (DMSO) or SHP099 (2 µM) for 6h, 24h, or 48h. **(B)** Statistical analysis (Student's t test) of cell cycle distribution data from **A**. *p<0.05, **p<0.01, ***p<0.001, ****p<0.0001.

A

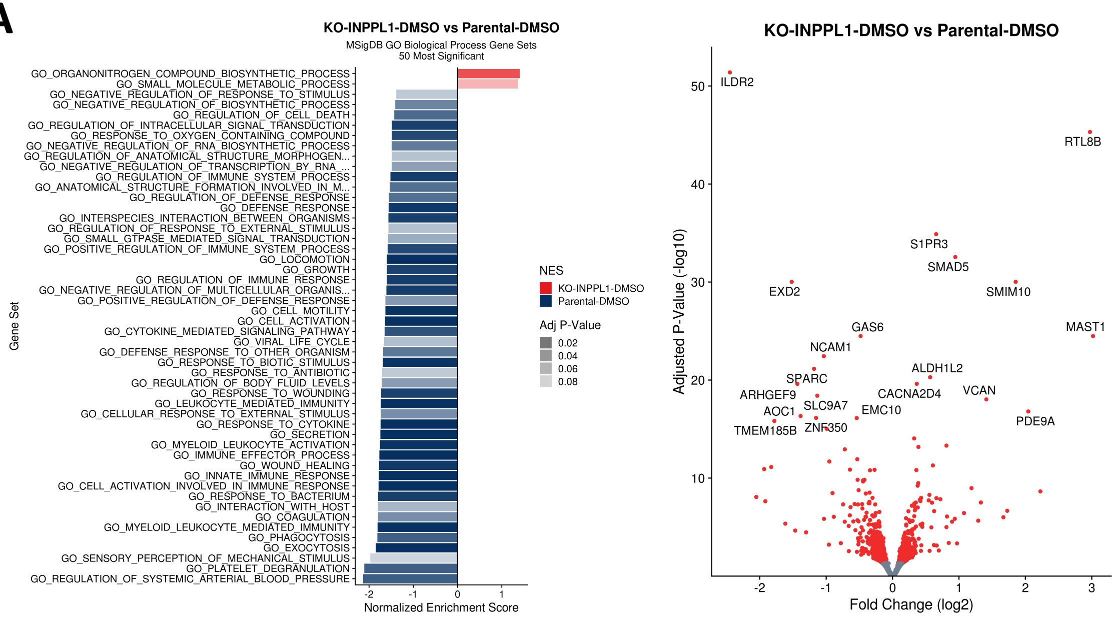


B

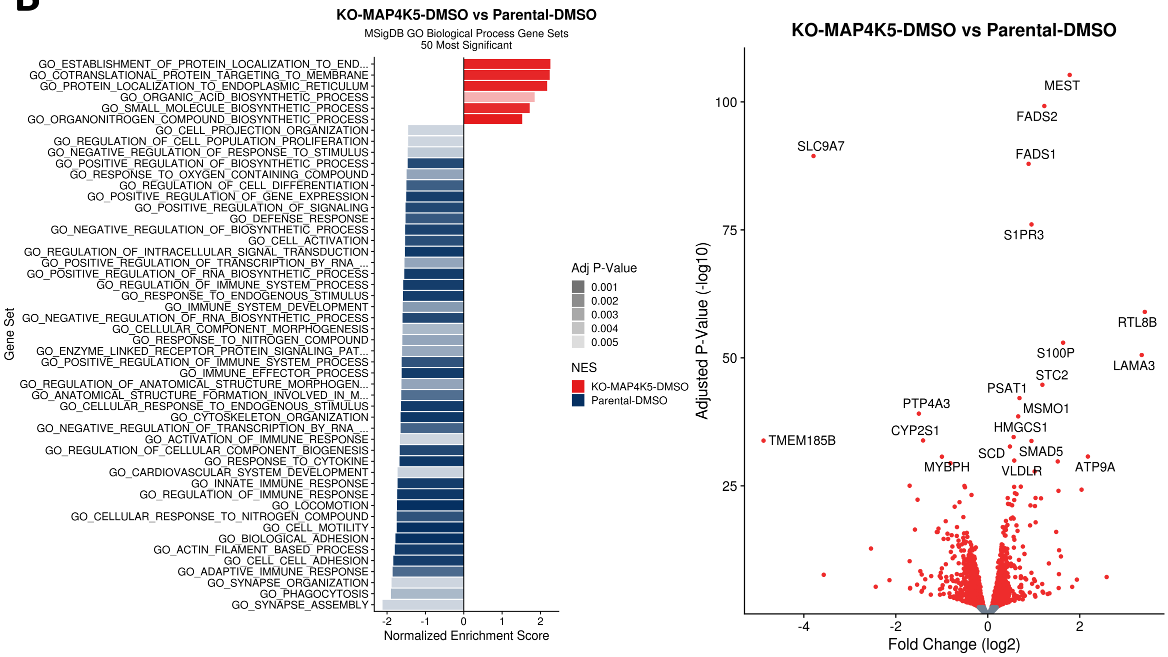


**Supplemental Figure 4. Effect of *INPPL1* or *MAP4K5* KO on basal gene expression in MOLM13 cells. (A)** Top 50 most significantly enriched MSigDB GO Biological Processes (left panel) and differentially expressed genes (right panel) in vehicle (DMSO)-treated *INPPL1* KO vs parental MOLM13 cells, as assessed by RNAseq. (B) Same as **A**, but comparing parental and *MAP4K5* KO MOLM13 cells.

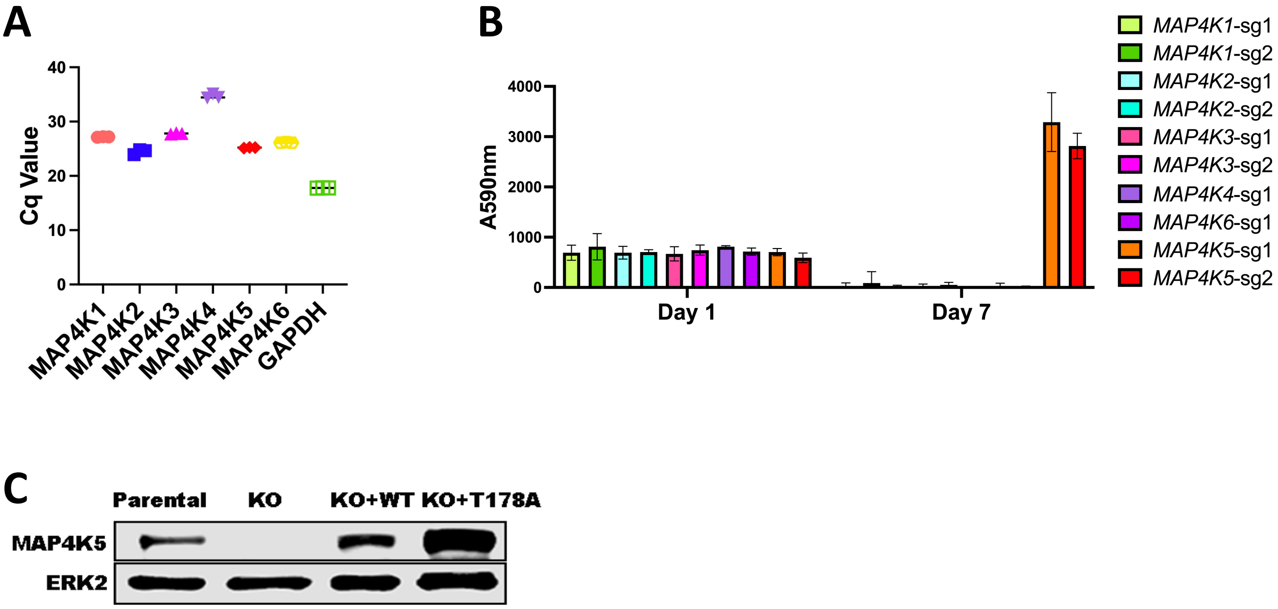


**Supplemental Figure 5. MAP4K5 kinase activity required to confer sensitivity to SHP099. (A)** All MAP4K family members are expressed at the RNA level in MOLM13 cells as detected by qRT-PCR. **(B)** Only *MAP4K5* KO causes resistance to SHP099. MOLM13 cells with deletion of the indicated *MAP4K* family member gene were treated with SHP099 (1 μM) and assessed by PrestoBlue assay at day 7. **(C)** Immunoblot showing reconstitution of *MAP4K5* KO MOLM13 cells with WT or kinase-dead (T178A) *MAP4K5*.

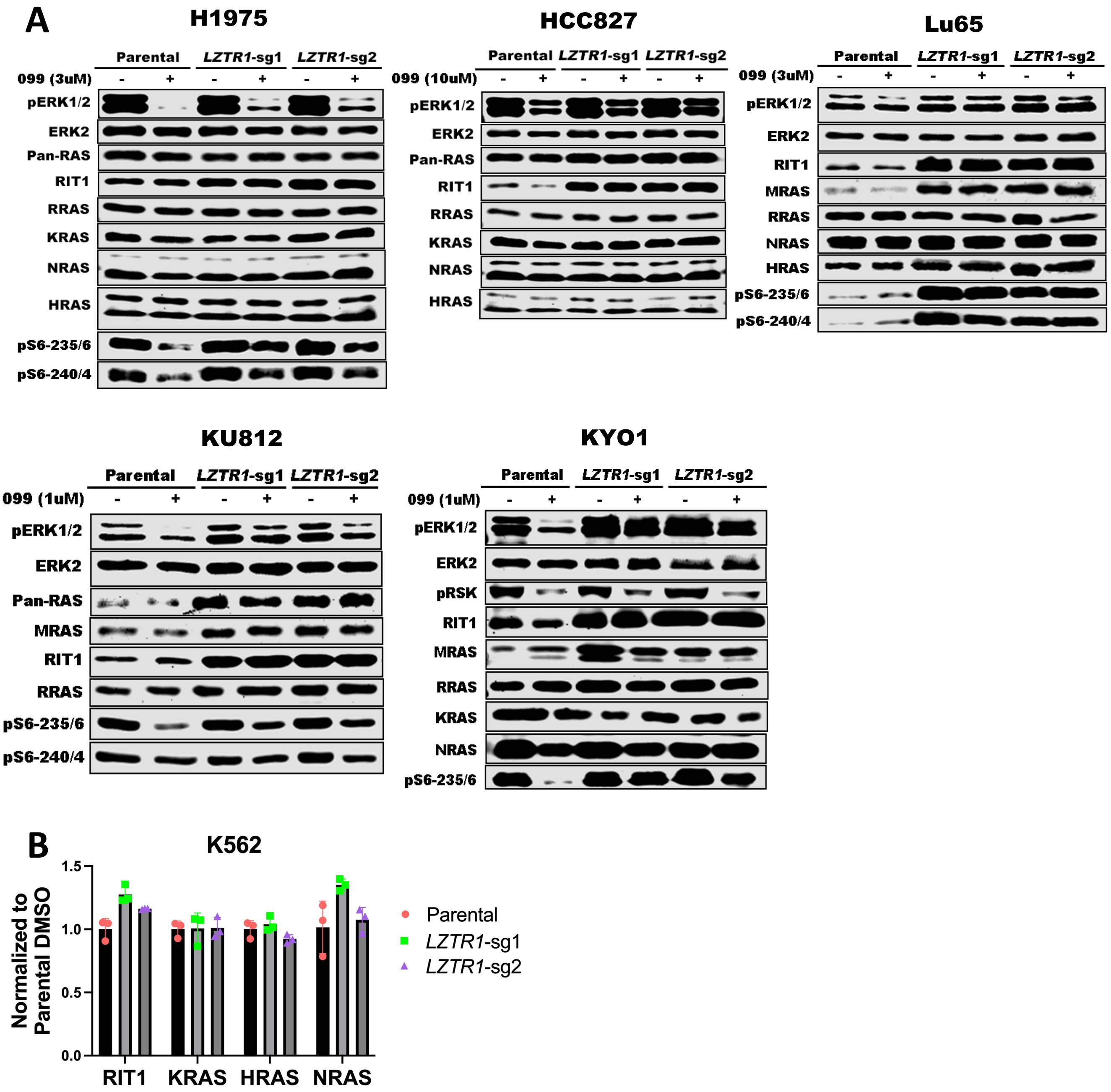


**Supplemental Figure 6. LZTR1 regulation of RAS family proteins is cell-context dependent. (A)** *LZTR1* KO differentially affects RAS family protein levels in H1975, HCC827, Lu65, KU812, and KYO1 cells. **(B)** RIT1, KRAS, HRAS, and NRAS mRNA levels in K562 cells are not affected by *LZTR1* KO.
